# TRIM33 loss reduces Androgen Receptor transcriptional output and H2BK120 ubiquitination

**DOI:** 10.1101/2025.02.14.638236

**Authors:** Nils Eickhoff, Janina Janetzko, Nuno Padrão, Sebastian Gregoricchio, Joseph C. Siefert, Liesbeth Hoekman, Simon Linder, Onno Bleijerveld, Andries M. Bergman, Wilbert Zwart

## Abstract

The Androgen Receptor (AR) is a ligand-dependent transcription factor that drives prostate cancer development and progression. Although, a detailed effect on AR biology has been described for a number of interacting proteins, many AR coregulators remain to be characterized in relation to their distinct impact on AR function. Here, we describe TRIM33 as a conserved AR-interactor across multiple prostate cancer cell lines. We observed that TRIM33 and AR share overall chromatin interaction profiles, in which TRIM33 is involved in downstream responsive transcriptomic output. In contrast to prior reports, we show that TRIM33 does not impact AR protein stability, but instead propose a model in which TRIM33 facilitates maximal AR activity by interfering with H2BK120 ubiquitination levels.

## INTRODUCTION

Activation and control of transcription factor (TF) activity is essential for cellular homeostasis and adequate response to extracellular stimuli ^1^. The activity of TFs can either be regulated indirectly by upstream signaling cascades (e.g., Wnt, β-Catenin ^2^) or directly through ligand binding ^3^. The latter group of TFs includes steroid hormone receptors, to which the estrogen, glucocorticoid and androgen receptor belong. The androgen receptor (AR) is a hormone-responsive transcription factor, that regulates the cellular response to the male sex hormone testosterone. It is well established, that the AR is involved in diseases like muscular atrophy ^4^, androgen insensitivity syndrome ^5^ and prostate cancer, in the latter being considered the key driver in development and progression of the disease ^6^. This causal role in prostate cancer renders AR the main target in prostate cancer therapy ^7,8^. Unfortunately, despite an initial response to AR-inhibition for most patients, eventual relapse to treatment is inevitable, as the cancer reactivates the AR signaling pathway despite low ligand levels ^9^. Therefore, a more in depth mechanistic understanding of the critical components involved in AR signaling is key to improve treatment ^10^.

On a molecular level, AR resides in the cytosol in absence of testosterone ^11^. Upon ligand binding, the receptor translocates to the nucleus, where it can bind DNA at so-called androgen receptor binding sites (ARBS), located mainly at putative enhancer elements. These are positive for enhancer-related histone marks (e.g., H3K4me1), EP300 and the active histone modification H3K27ac ^12^. At these sites, AR harbors *cis*-regulatory activity to drive expression of its target genes. To facilitate alterations in gene expression, AR interacts with numerous proteins to form its canonical transcription complex and recruit the epigenetic machinery to alter DNA accessibility and local epigenetic state to finally alter transcriptional output ^13^.

Recent technological advances provided tools that allow for a systematic identification of AR interacting proteins in a comprehensive unbiased fashion. These technologies, such as RIME (Rapid Immunoprecipitation and Mass spectrometry of Endogenous proteins; ^14^), ChIP-SICAP (Chromatin Immunoprecipitation coupled with Selective Isolation of Chromatin-Associated Proteins; ^15^) and Bio-ID ^16^ identified both known and well-established AR interacting proteins as well as previously unknown AR-interacting proteins. Among these novel AR interactors are several components of the transcription intermediary factor 1 (TIF1) family ^17–19^, which consists of 4 proteins (TRIM24 – TIF1α, TRIM28 – TIF1β or KAP1, TRIM33 – TIF1γ, TRIM66 – TIF1δ) that belong to the tripartite motif (TRIM) containing protein family ^20^. Interestingly, TRIM24, TRIM33 and TRIM66 contain nuclear receptor interacting motifs ^21,22^ and have all been associated with prostate cancer development ^23^. What makes these proteins especially interesting in the light of transcriptional regulation is the fact that they contain both a Bromo and a PHD domain at their N-terminus ^20^. Both domains are epigenetic readers of histone modifications allowing to potentially integrate two histone marks on one histone tail by one reader protein ^24^.

Here, we analyzed the AR protein interactome using RIME across six different prostate cancer cell line models and defined a core AR interactome including TRIM33. To study the role of TRIM33 in AR biology, we generated genomics and proteomics datasets of TRIM33 wildtype and knockout cell lines. We show here, that TRIM33 and AR co-occupy most of their genomic binding sites and TRIM33 loss altered expression of a subset of AR-responsive genes. In contrast to prior studies, AR levels were not affected by TRIM33 loss, and despite the canonical E3 ligase annotation of TRIM33, no indication of altered protein degradation was observed upon TRIM33 loss. Instead, we observed that TRIM33 expression coincided with reduced H2BK120 ubiquitination at genes under shared AR/TRIM33 control; a histone modification implicated in transcriptional regulation and higher order chromatin organization ^25^. Altogether, we propose an alternative model of how TRIM33 impacts AR signaling, independent of protein degradation ^26^.

## RESULTS

### TRIM33 is part of a core AR interactome across prostate cancer cell lines

To systematically characterize proteins that interact with AR across various prostate cancer disease stages, we performed AR RIME experiments in AR expressing prostate cancer cell lines that were sensitive (LNCaP, LAPC4) or resistant (CWR-R1, 22Rv1, LNCaP-abl, 42D) to hormone deprivation. This allowed us to model disease progression from a hormone-sensitive state to castration resistance. An AR negative cell line (PC3) was used as a negative control. In PCA space, immunoprecipitations (Antibodies) of the IgG negative control separated clearly from the AR ones (**Figure 1A**), as well as the AR positive cell lines from the AR-negative PC3 cells (**Supplementary Figure 1A**). After filtering for nuclear proteins, the AR positive cell lines showed an enrichment of around 400 proteins (LNCaP = 423, LAPC4 = 304, 22Rv1 = 343, CWRR1 = 408, 42D = 433, LNCaP-abl = 385; PC3 = 35) as potential AR interactors (**Supplementary Figure 1B-H**), of which 119 were shared between all (**Figure 1B**, **Supplementary Table 1**), hereafter referred to as the “AR core interactome”. Over-representation analysis of these proteins against the CORUM database of protein complexes showed enrichment for complexes involved in epigenetic regulation such as the SWI/SNF (BAF) complex, involved in nucleosome positioning, or the histone demethylase LSD1, both of which are associated with AR biology ^27,28^ (**Figure 1C**). Interestingly, TRIM24 and TRIM33 were found in the AR core interactome. TRIM24 has been extensively studied in relation to AR biology ^17,29^, but the impact of TRIM33 on AR is less well understood. To identify which proteins may be part of a shared TRIM33-AR complex, we next performed TRIM33 RIME in LNCaP cells with AR stimulation (5nM R1881, 4h) and observed enrichment of 185 nuclear proteins as potential TRIM33 interactors (**Figure 1D**), of which 19 were also part of the AR core interactome (**Figure 1E**). The majority of these proteins were associated with canonical AR transcriptional function including FOXA1 ^30^, HOXB13 ^31^, NKX3-1 ^32^ and the SWI/SNF complex ^27^. Similarly, we charted the TRIM24 interactome and we could confirm AR interaction alongside interaction with SWI/SNF members and TRIM33 (**Supplementary Figure 1I**). Overall, the overlap between enriched TRIM33 and TRIM24 interactors was 49 proteins including AR, FOXA1, SWI/SNF complex members and RNF20/40.

**Figure 1.**
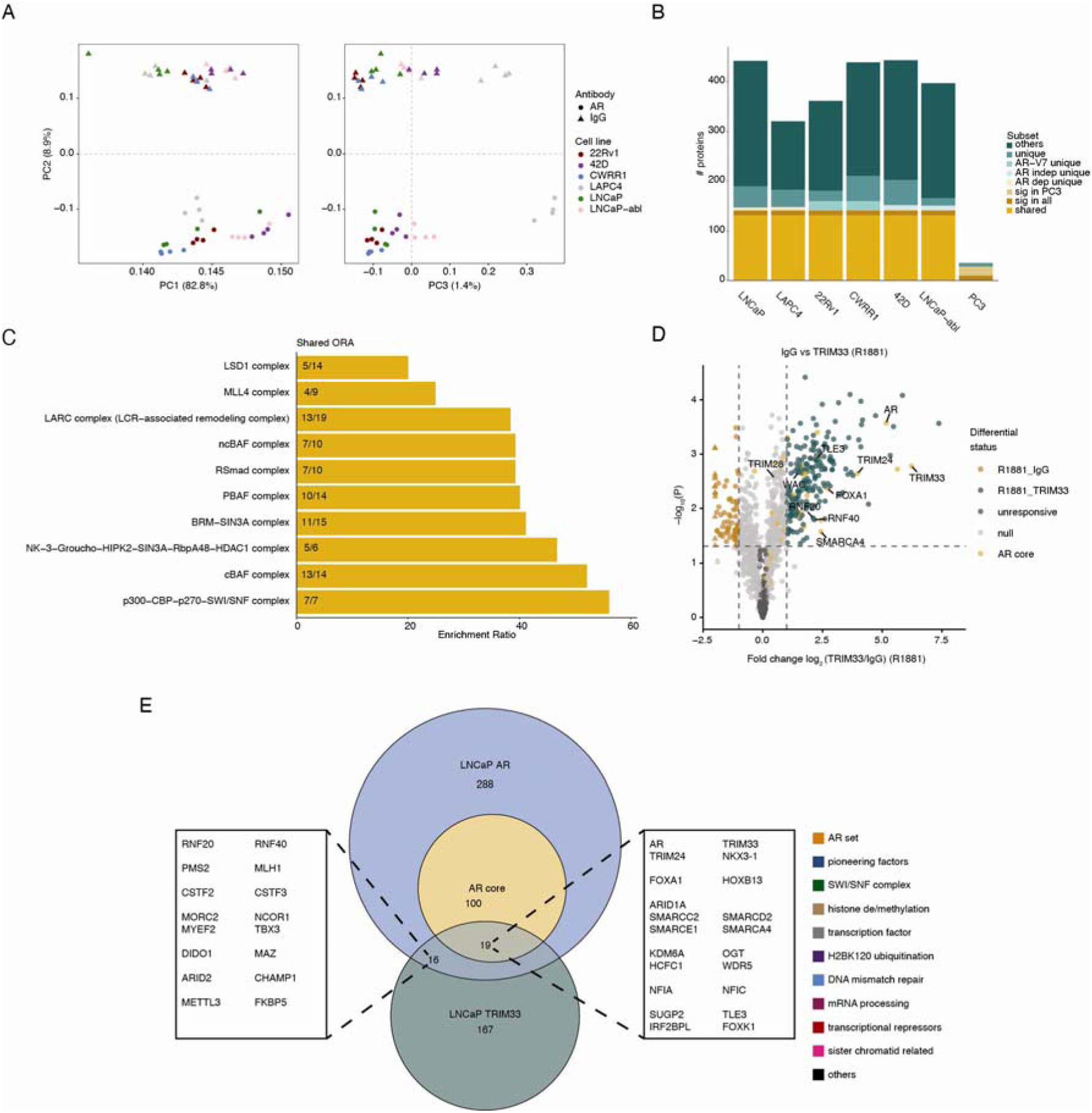
TRIM33 is part of the AR core interactome. **A** PCA of AR and IgG RIME data in AR positive cell lines. **B** Barplot representing the enriched nuclear proteins (LFQ difference > 1 & p-value < 0.05) per cell line assigned to the displayed subsets. Shared = found in all AR positive cell lines and not in PC3. Sig in all = found across all cell lines. Sig in PC3 = proteins significant in PC3 and another cell line. AR dep unique = Shared between LNCaP, LAPC4 and no other. AR indep unique = shared between CWR-R1, 22Rv1, LNCaP-abl and 42D and no other. AR-V7 unique = proteins shared in cell lines that express the alternative AR splive variant AR-V7 (CWR-R1, 22Rv1, **Supplementary Figure 3A**) **C** Over representation analysis of the AR core proteins against the CORUM protein complex database **D** Volcano plot displaying the TRIM33 RIME data with the AR core proteins highlighted in orange. Triangles depict proteins enriched in IgG with fold changes out of the axis limits. **E** Euler Diagram of the AR core, LNCaP AR and TRIM33 RIME nuclear enriched proteins.

### TRIM33 and AR share chromatin binding profiles

To study if the AR/TRIM33 and AR/TRIM24 interaction is taking place on chromatin and how this may influence the epigenetic landscape, we performed ChIP-seq experiments for TRIM33 and TRIM24 and two histone marks that have been associated with the Bromo (H3K18ac) and PHD (H3K9me3) domains of TRIM33 ^33–35^, in hormone-deprived conditions (DMSO) and two timepoints after stimulation with the synthetic androgen R1881 (4h and 24h) (**Supplementary Figure 2A-C**).

The majority of TRIM33 peaks (DMSO = 1695, 4h = 1672, 24h = 5984, averages across replicates, see **Supplementary Figure 2C**), across all timepoints, were found in distal intergenic and intronic genomic regions, suggestive of binding to *cis*-regulatory enhancer elements, which is similar to binding patterns previously reported for AR (**Figure 2A**, GSE94682 (Stelloo et al., 2018)). Interestingly, the proportion of promoter overlapping peaks for TRIM33 was the highest in hormone-deprived DMSO conditions. Next, we linked peaks to the closest gene and performed over representation analysis (ORA) against the cancer hallmark gene sets. The stimulated conditions showed enrichment for the androgen response pathway, suggesting that TRIM33 plays a role in the regulation of these core AR target genes (**Figure 2B**). When overlapping the TRIM33 peaks with AR binding upon 4h stimulation, most TRIM33 binding sites were shared with AR (2896; 89%), with merely 356 peaks being only present for TRIM33 (**Figure 2C**). Both sets showed clear induction of TRIM33 binding upon AR stimulation, and an increase in H3K18ac signal at these sites. Interestingly, H3K18ac was also found elevated at AR peaks that did not contain TRIM33 called peaks after stimulation. This may possibly be explained by the observation that TRIM33 signal at these sites increased upon AR stimulation but not enough to pass peak calling thresholds. For H3K9me3 however, its broad distribution across repressed heterochromatin regions ^37^, did not allow for efficient peak calling. Nonetheless, TRIM33 peaks showed depletion of this mark at the peak center (**Figure 2C**). Lastly, TRIM33 chromatin patterns were virtually completely shared with TRIM24 (**Figure 2C-D**); another member of the TIF1 complex. These data indicate a possible formation of heteromeric TRIM complexes as suggested before ^38,39^. Apart from peak occurrence, we also investigated the dynamic changes in TRIM33 binding upon AR activation, through identification of peaks that changed significantly in intensity between conditions. This revealed 3248 sites for TRIM33 that were gained after 4h of stimulation compared to the hormone-deprived condition and 462 sites after 24h that were not yet significantly increased at 4h (**Figure 2D-E**, Supplementary Figure 2D). Interestingly, a distinct set of 164 TRIM33 binding sites was lost after 24h of stimulation compared to the unstimulated control. This loss coincided with reduced H3K18ac, absence of AR binding and the spreading of H3K9me3 across the peak (**Figure 2D-E**). Additionally, the TRIM24 signal already diminished upon 4h stimulation for TRIM33 lost sites after 24h stimulation (**Figure 2D-E**). To asses which other proteins might be co-occupying these binding sites, we overlaid the TRIM33 binding site subsets with those identified in publicly available ChIP-seq data sets (n= 13,976) as part of the Cistrome Data Browser TF ChIP-seq sample collection ^40^ (**Supplementary Figure 2F**). For the 4h TRIM33 gained sites, we found enrichment of AR and its interactors (e.g., SMARCA4, FOXA1, ARID1A; **Figure 2F**). This further confirms, that TRIM33 and AR share distinct genomic binding sites and that TRIM33 recruitment to the DNA is enhanced by active AR signaling.

**Figure 2.**
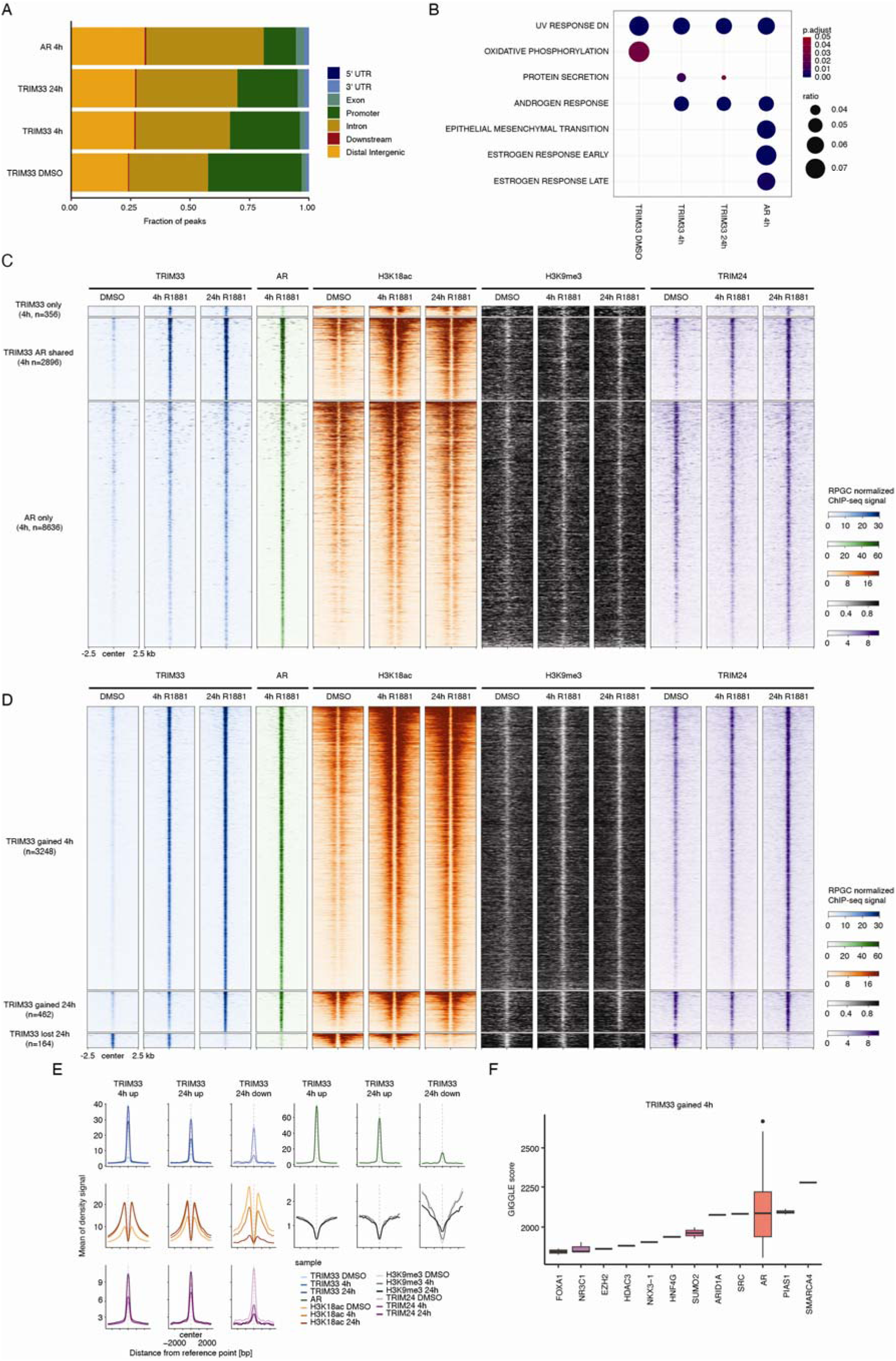
TRIM33 and AR co-occupy similar genomic locations. **A** Feature distribution of peaks identified in the given ChIP-seq experiment. For AR, ChIP-seq data from GSE94682 was used **B** Over representation analysis of peak associated genes. Shown are only genesets that have a p.adjust < 0.05. **C** Tornado plot for the TRIM33 and AR shared and unique peaks at 4h stimulation. Data represents the average of QC passing replicates. **D** Tornado plot for the TRIM33 sites with dynamic behavior. Data represents the average of QC passing replicates. **E** Average signal intensity plots for the regions shown in D. Lighter colors represent the DMSO control whereas darker colors represent stimulated conditions with R1881. **F** GIGGLE results for the TRIM33 sites gained after 4h of R1881 treatment

### TRIM33 KO reduces transcription of AR target genes

To investigate the functional impact of TRIM33 on AR action, we generated CRISPR/Cas9 mediated knockouts for TRIM33 in LNCaP prostate cancer cells (**Figure 3A**). In an initial analysis, we investigated whether TRIM33 influences prostate cancer cell proliferation. However, no significant or consistent differences were observed in monoclonal TRIM33-knockout lines (**Figure 3B**) or polyclonal TRIM33 knockouts across LNCaP and several other prostate cancer models (**Supplementary Figure 3A-E**), indicating that TRIM33 loss does not affect the proliferative potential of prostate cancer cells. Apart from its role in cancer cell proliferation, AR signaling also impacts other phenotypes, including protein secretion and cellular differentiation during development (Vickman et al., 2020). To investigate whether TRIM33 is involved in these signaling axes, we explored the transcriptomic alterations upon TRIM33 perturbations, by conducting RNA-seq experiments in two different monoclonal knockout cell lines (clones C2 and F2), the polyclonal knockout in LNCaP and 42D cells under hormone-deprived conditions and upon R1881-mediated AR activation. On PCA space, the two cell lines separated clearly, as well as the R1881 treated samples from the unstimulated control (**Supplementary Figure 4A**). For LNCaP alone there was separation along PC1 by treatment but there was also separation along PC2 which is driven by the F2 clone (**Supplementary Figure 4B**). In the non-targeting control (NT), 265 differentially expressed genes (DEGs) were detected upon 6h R1881 treatment (**Supplementary Figure 4C**). This number of treatment specific DEGs was reduced in both knockouts with 201 for F2 and 120 for C2, of which the majority overlapped with the DEGs of the control following R1181 treatment (F2: 189, C2: 108; **Figure 3C**). The magnitude of decrease in AR response is in line with residual TRIM33 levels, showing the least effect in the polyclonal knockout and the most-profound effect in the monoclonal C2 knockout cell line (**Figure 3D**). When comparing the stimulated conditions between control and clone C2, we observed that key AR target genes, like *KLK3*, were decreased in their expression, along with *TRIM33* itself (**Figure 3D-E**). Comparable observations were made for the second clone F2 (**Supplementary Figure 5A-B**). Gene set enrichment analysis (GSEA) against the cancer hallmark gene sets, showed that androgen response was downregulated in both clones, compared to the non-targeting control (**Figure 3F** and **Supplementary Figure 5C-D**). To investigate the potential of TRIM33 as a transcriptional regulator, we integrated the ChIP-seq and RNA-seq data streams. This analysis revealed that TRIM33 has the potential to regulate the expression of several AR responsive genes (e.g., *NKX3-1*, *FKBP5* or *STEAP4*), as well as a group of downregulated genes that do not belong to the AR signaling axis (**Figure 3G**).

**Figure 3.**
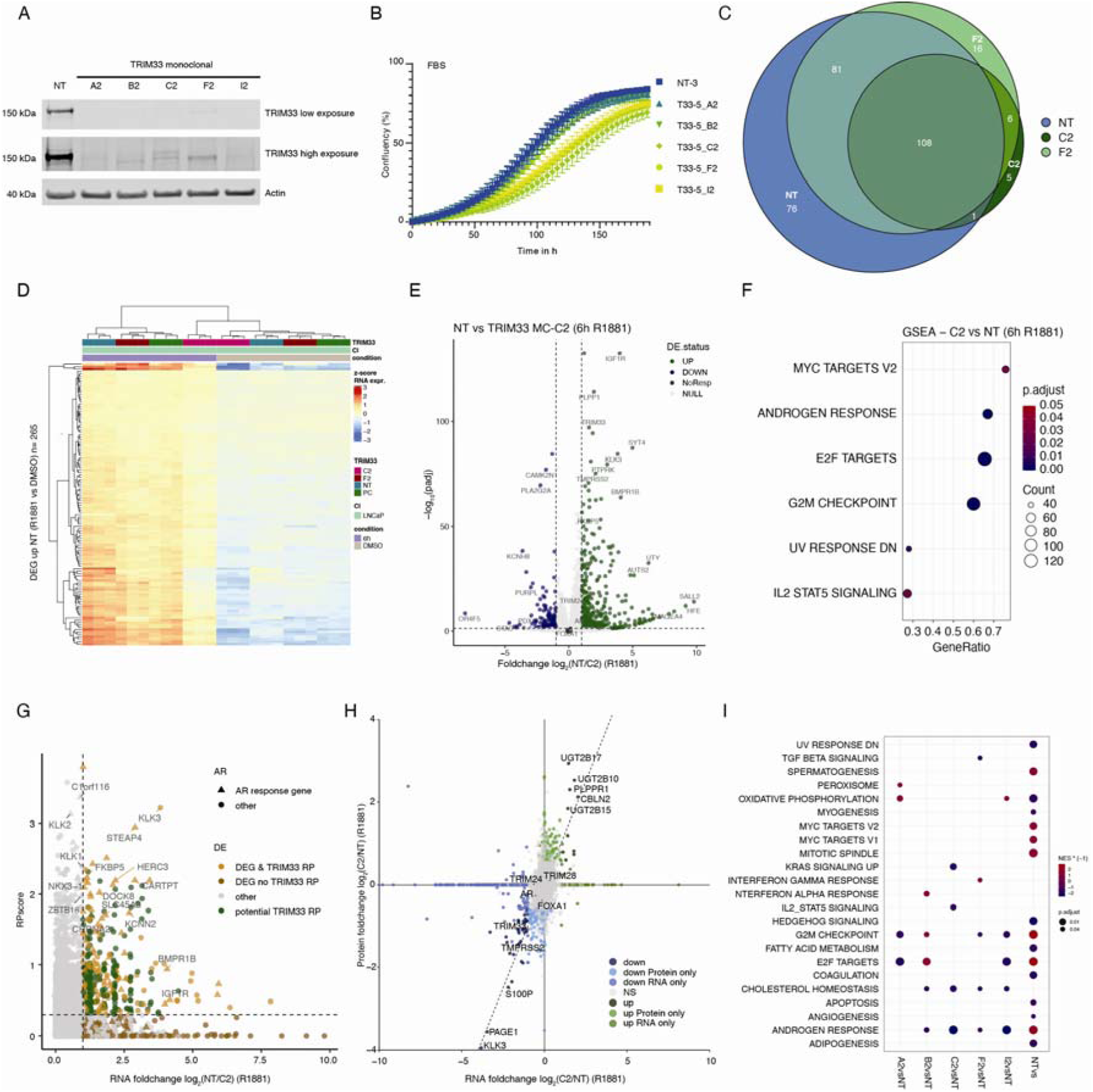
TRIM33 loss reduces AR transcriptional output. **A** Western blot for TRIM33 across the generated monoclonal knockout cell lines of the TRIM33-5 guide. **B** Incucyte growth curves of monoclonal TRIM33 knockout cell lines in FBS. As an error SEM is shown. **C** Euler diagram for all DEGs in the represented cell lines between DMSO and 6h R1881 stimulation. **D** Heatmap for DEGs in the non-targeting control across all sequenced LNCaP cell lines. PC represents the TRIM33 polyclonal knockout parental cell line. **E** Volcano plot of gene expression levels between TRIM33 monoclonal knockout C2 and the non-targeting control at 6h stimulation with R1881. Genes in green are higher expressed in non-targeting cells than knockout cells. **F** GSEA of the data in E. Shown are only genesets that have a p.adjust < 0.05. **G** Scatterplot of the Cistrome-Go analysis. Used are the TRIM33 peaks at 4h stimulation and the RNA foldchange showed in E. Triangles depict AR response genes as defined by differential expression status in non-targeting control with and without AR stimulation. Cut-off for regulatory potential was set to 0.3. **H** Scatterplot comparing TRIM33 knockout (C2) to non-targeting foldchanges on RNA (x-axis) and protein (y-axis) levels. Genes or proteins just found in one dataset were set to 0 in the other. The dashed line represents the diagonal with a slope of 1. Cut-offs for classification were a foldchange > 1 for RNA and > 0.5 for proteomic data. **I** GSEA of proteomic data of all tested TRIM33 knockouts in R1881 treated conditions compared to the non-targeting. NTvs represents the comparison of non-targeting cells with and without stimulation.

As TRIM proteins, including TRIM24 ^42,43^ and TRIM33 ^26,44^, have been implicated as E3 ubiquitin ligases, we next explored the effect of TRIM33 loss on the proteome of LNCaP cells. Comparing these proteomics data with our transcriptomics data revealed overall a very concordant correlation between both techniques (**Figure 3H**). Performing GSEA on the proteomics data showed that only androgen response and cholesterol homeostasis were downregulated in all knockouts, relative to the non-targeting control (**Figure 3I**). Jointly, these data suggest that the observed effects are mainly driven by transcriptional changes due to TRIM33 loss rather than post-translational protein degradation. Additionally, we do not observe changes of AR protein levels, which is in contrast to prior work that linked TRIM33 and AR degradation via SKP2 ^26^ (**Supplementary Figure 5E)**.

Lastly, we analyzed whether TRIM33 absence would affect the AR interactome, but no differences in the AR interactome were observed between KO and non-targeting controls, except for TRIM33 itself (**Supplementary Figure 5F**).

### TRIM33 loss reduces H2BK120ub at AR target genes

As TRIM33 is the reader of H3K18ac, we next checked if loss of TRIM33 affected H3K18ac stability or AR binding to the chromatin. Therefore, we performed ChIP-seq for AR and H3K18ac in the TRIM33 knockout and non-targeting cell lines (**Figure 4**). Additionally, we took along H2BK120 ubiquitination (H2Bub), a marker for active transcription across gene bodies ^25^, as it has been reported that TRIM33 and TRIM24 work in concert to facilitate this mark at the *HSP72* promoter together with HSF1 in HeLa cells ^38^. In PCA space, the different IP targets (AR, H3K18ac, H2Bub) formed distinct clusters (**Figure 4A** top left). Individual analysis for each target showed no clear separation between knockout and control samples. Instead, the primary distinction was between untreated and R1881-stimulated samples (**Figure 4A**). For H2Bub the least differences could be observed in PCA space, which could be explained by the fact that the majority of gene bodies throughout the genome are not affected by TRIM33 loss and a more focused approach is needed (**Figure 4A**, Supplementary Figure 6A-B). Differential binding analysis between TRIM33-KO and control across all timepoints showed no differences for H2Bub and a small fraction of peaks changing for both H3K18ac and AR (**Supplementary Figure 6C-D**). Density profiles at the small subset of TRIM33-KO affected peaks for AR and H3K18ac showed weak binding and noisy behavior, suggesting that these might be artifacts (**Supplementary Figure 6C-D**, top panels). Across the identified differential binding sites for TRIM33 after stimulation, the signal intensities were relatively similar across all subsets (**Figure 4B**). We then tested whether TRIM33-KO affected H2Bub signal for genes that are differentially expressed upon TRIM33 perturbation, as determined by RNA-seq (**Figure 3D**). Upregulated genes in TRIM33-KO cells had higher levels of H2Bub compared to non-targeting control (**Figure 4C-D**). Interestingly, AR stimulation reduced H2Bub at these sites in both the knockout as well as the non-targeting compared to the unstimulated condition. For the genes downregulated upon TRIM33 knockout, the opposite behavior was observed, with higher H2Bub in the non-targeting as compared to the knockout cell line. Statistical analyses (with a p-value cut-off of 0.01) showed that only the differentially expressed genes are affected on H2Bub levels whereas the H2Bub signal for unresponsive genes was not altered (**Figure 4E**).

**Figure 4.**
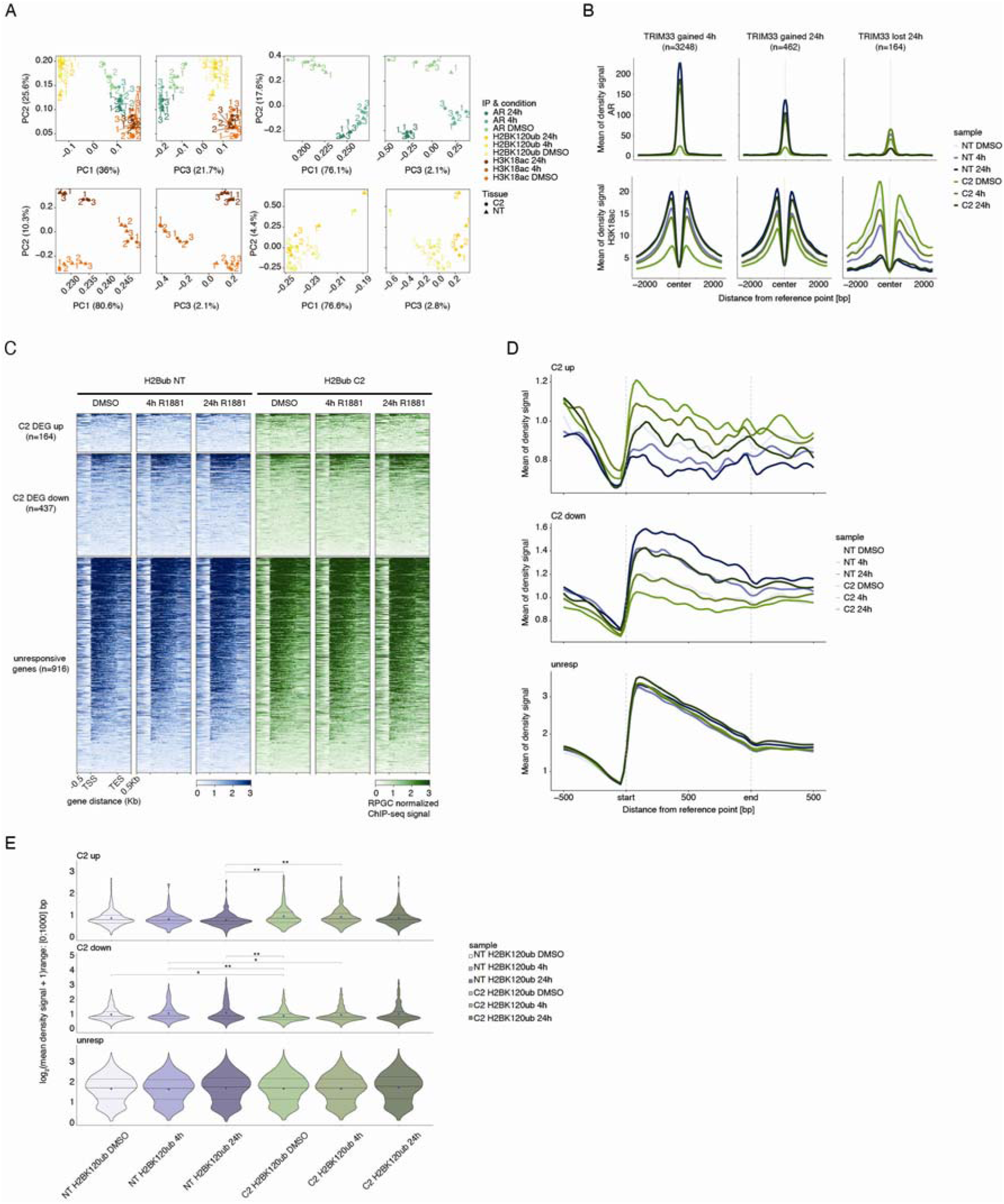
AR target genes have lower H2Bub levels in TRIM33 knockout cells. **A** PCA plots for grouped and individual analysis of the TRIM33 C2 knockout ChIP-seq data. **B** Average signal intensity profiles of AR (top) and H3K18ac (bottom) across the tested cell lines on the TRIM33 R1881 treatment gained and lost regions of Figure 2D. TRIM33 C2 knockout cells are in green and non-targeting in blue. **C** Heatmap of H2Bub signal across gene bodies of differentially expressed genes from Figure 3E as well as the unresponsive genes. **D** Average signal intensity plots for the regions depicted in C. **E** Distribution of the sum of the H2Bub signals across the scaled gene body from the regions in C. Significance levels of Wilcoxon ranked sum test: * < 0.01, ** < 0.001

Together these data support the notion that TRIM33 is neither affecting AR chromatin occupancy nor AR turnover, but instead impacts the transcriptional output of AR target genes (Supplementary Figure7A-C). In this role, TRIM33 is required for full AR activity, which is tightly associated with distinct alterations in H2BK120 ubiquitination in a locus-specific manner.

## DISCUSSION

Despite advances in better understanding the AR cistrome throughout the development, treatment and progression of prostate cancer (Linder et al., 2022; Pomerantz et al., 2015; Stelloo et al., 2018), the effect of non-DNA binding cofactors on AR biology remains incompletely understood. Furthermore, while it is established that distinct AR-subcomplexes form on a genome-wide scale (Stelloo et al., 2018), the implications thereof remain understudied. Here we characterized TRIM33 in the light of AR biology, and showed that TRIM33 is required for complete transcriptional output of AR at distinct genomic locations. These changes are associated with alterations in H2BK120 ubiquitination status at TRIM33/AR-coregulated genes. This finding is in contrast to the previously reported role of TRIM33 on AR biology, that reported TRIM33 levels to affect AR protein levels via SKP2 ^26^. These discrepancies could be due to the different depletion methodologies and timeframes, as the authors of the prior report used transient siRNAs for depletion, while the current study made use of stable CRISPR/Cas9 mediated knockout models. To investigate whether these discrepancies were due to the different methodologies in TRIM33 depletion, we carried out experiments employing both the TRIM33 siRNAs described in the prior study ^26^ as well as the commercially available siRNA SMARTpool (Dharmacon). Surprisingly, while in our hands only the SMARTpool showed on-target depletion of TRIM33, none of the siRNAs had any effect on AR protein levels (**Supplementary Figure 8A**). Further investigations would be needed to resolve these inconsistencies between the two studies.

TRIM24, TRIM28 and TRIM33 are all members of the TIF1 complex, and while prior studies have investigated the relative contributions of TRIM24 and TRIM28 on AR biology, TRIM33 remains largely understudied. While TRIM33 is classically annotated as a E3 ligase, our data suggest that TRIM33 mostly acts on the transcriptional level, rather than altering the proteome through post-translational ubiquitination. It is important to consider that not all ubiquitin modifications lead to proteasomal degradation of the ubiquitinated protein ^47^. Therefore it is possible that TRIM33 is involved in other ubiquitin linkages, like it has been shown for TRIM24 ^43^ or mono-ubiquitination as is suggested for H2BK120. For the latter, even though recruited individually, prior work reported that H2Bub was only deposited when both TRIM24 and TRIM33 were present ^38^. Our study cannot clearly decipher whether TRIM33 is actively influencing H2Bub or whether the altered H2B ubiquitination is a consequence of reduced transcription through TRIM33 action on AR enhancers. It is also possible that TRIM33 ubiquitinates other proteins in the AR complex, altering their action in a proteasome independent fashion. Implementation of ubiquitin-specific probes may shed light on these possibilities.

Notably, TRIM33 plays a role in transcription regulation at merely a subset of AR sites. These observations highlight that cofactors are not universal at all AR binding sites, leading to subcomplexes owing distinct functions (Stelloo et al., 2018). In our study, while we observed that proliferation is not affected by TRIM33 knockout, hallmark AR target genes are nonetheless downregulated. These data implicate that the hallmark AR gene set does not reflect the role of AR in proliferation, but rather is indicative for its role in other cellular functions. To better capture the oncogenic roles of AR, it may proof critical to rather select genes affected in clinical transitions in tumorigenesis or metastasis formation instead ^48^.

Current technologies for AR protein interactomes are averaged across all binding sites, which leads to a loss of dependencies and co-occurrences between proteins in the AR complex. Single locus transcription factor complex analyses have been shown to be possible, but remain technically highly challenging ^49^. Alternatively, overlaps of interactomes of several AR interactors can shed light on co-occurrence patterns, as we can see here for the 35 proteins shared between AR and TRIM33 in LNCaP cells. In general, the interactome datasets between orthogonal measures appears surprisingly small, highlighting the necessity of stringent controls and analysis to increase true positive hits.

All together, we confirm the action of TRIM33 as a coactivator of AR, however not via AR stability but rather through attenuating maximal transcriptional activity potentially by altering H2Bub levels in a locus-specific manner. Even though not impacting prostate cancer cell proliferation in our hands, TRIM33 appears crucial for the expression of canonical AR target genes.

## METHODS

### Cell culture

LNCaP, 22Rv1, CWRR1, PC3, LAPC4 and HEK293T were purchased from ATCC and LNCaP-abl, 42D were a generous gift from Amina Zoubeidi (Vancouver). Prostate cancer cell lines were cultured in RPMI 1640 medium (Gibco, Thermo Fisher Scientific) supplemented with Penicillin/Streptomycin (Gibco, Thermo Fisher Scientific; 1U, 1µg/ml) and 10% FBS (LNCaP, 22Rv1, CWR-R1, LAPC4; Capricorn) or 5% of dextran and charcoal stripped FBS (DCC, LNCaP-abl). 42D cells were continuously cultured in the presence of 10µM Enzalutamide (MedChemExpress). HEK293T cells were cultured in DMEM medium (Gibco, Thermo Fisher Scientific) supplemented with Penicillin Streptomycin and 10%FBS. For experiments involving hormone stimulation, cells were cultured in DCC for 72h prior to the start of the experiment. Cells were regularly tested for *Mycoplasma* and were authenticated by short tandem profiling (Eurofins Genomics).

### Knockout generation

Guides targeting TRIM33 (T33-3: 5’-ACAGAGTCTGTTGGAGCATC-3, T33-4: 5’-ACTATGGCAAATGCAAACCG-3, T33-5: 5’-CTCCTCCTCCACCAGCACCG-3) or non-targeting control (NT: 5’-AACTACAAGTAAAAGTATCG-3) were cloned into lentiCRISPRv2 plasmids (^50^ Addgene: #52961). For lentivirus production, CRISPR plasmids were cotransfected with 3^rd^ generation lentiviral plasmids (5:1:1:1) using polyethylenimine (Polysciences) into HEK293T cells. Virus containing supernatants were harvested, filtered using a 0.22µm anti-pyrogenic filter, and stored at -80°C until use. Two days post transduction, cells were selected with puromycin (Sigma-Aldrich, 2µg/ml) for 2 weeks and knockout efficiencies were estimated using western blotting. For monoclonal cell lines, polyclonal parental cells were FACS sorted into 96 well plates (Greiner, Cellstar) and gradually grown out.

### Western blot

Whole cell lysates were prepared using a 2x Laemmli lysis buffer supplemented with EDTA-free complete protease inhibitor cocktail (Roche), 100mM NaF (Thermo Scientific) and 2mM Na_3_VO_4_ (Jena Bioscience). Lysates were sonicated for 10 cycles 1 second on/off with a probe sonicator (Active Motif) at 20% amplitude. Protein amounts were measured using the Pierce BCA protein assay kit (Thermo Fisher Scientific) according to the manufacturer’s instructions. Then 30µg of protein were reduced using 0.1M DTT (Sigma Aldrich) and loaded onto 4-12% Bis-Tris NuPAGE gels, run in MOPS buffer and subsequently wet transferred onto 0.45µm nitrocellulose membranes (Santa Cruz Biotechnology). Membranes were blocked in 5% fat-free milk in PBS-0.001% Tween20 (Sigma Aldrich) prior to overnight incubation at 4°C with primary antibodies (Actin: MAB1501R, Merck, 1:3500; TRIM33: D7U4F, Cell Signaling Technologies, 1:1000, 1:1000; AR: 06-680, Merck, 1:1000). Subsequently, the membranes were incubated with appropriate IRDye® secondary antibodies (680 or 800nm, LI-COR, 1:10,000). Signal detection was carried out using an Odyssey CLx imager (LI-COR) and image analysis was performed using Image Studio Lite (v5.5)

### Cell proliferation

#### Incucyte

1000 LNCaP cells were seeded in 384-well plates in DMSO or 10µM Enzalutamide (MedChemExpress) containing medium. After 24h the plate was transferred into an Incucyte® ZOOM live cell analysis system (Sartorius) and wells were imaged every 4h over the course of the experiment. Confluency was calculated with the build-in software with an adjusted mask (Segmentation adjustment = 0.1, Hole fill ≤ 40 µm^2^, Area ≥ 500 µm^2^).

#### CellTiter-Glo

CellTiter-Glo (Promega) assays were used as an orthogonal method to measure cell viability. Cells were seeded into 96-well plates (LNCaP: 750, LAPC4: 1000, 42D: 1000, CWRR1: 500; all cells/well). After 24h cells were treated either with DMSO or 10µM Enzalutamide and, after 7 days, signal was measured according to the manufacturers protocol and analyzed using GraphPad Prism (9.4.1).

### Whole cell proteomics

For whole cell proteomics, 700,000 LNCaP cells were seeded in a 6cm-dish. 48h later the cells were washed twice with cold PBS on ice and then scraped twice in 500µL PBS. Cells were pelleted at 1000 rpm at 4°C for 5 min. The supernatant was removed and the pellets frozen in liquid nitrogen, and stored at -80°C until further processing.

Details of the mass spectrometry can be found in the **Supplementary Methods.**

### RIME experiments

Six confluent 15cm-dishes of a cell line cultured in DCC for 72h were induced with 5nM R1881 or DMSO for 4h before fixing with 1% formaldehyde (Electron Microscopy Sciences) for 10 minutes. Samples were processed as described in Mohammed *et al.* In brief, cells were harvested and chromatin was sonicated with a Bioruptor pico (Diagenode) sonicator using cycles of 30s on/off until obtaining a fragment sizes around between 250-700bp. 10µg of AR, TRIM24 or TRIM33 antibody were coupled to 50µL protein A dynabeads (Invitrogen, cat. 10008D) for immunoprecipitation. After over-night immunoprecipitation, beads were washed with RIPA-RIME buffer and 100mM ammonium bicarbonate and stored at -80°C until on bead tryptic digest was performed and protein content quantified.

#### Mass spectrometry

For mass spectrometry, peptide mixtures were prepared and measured as previously described (Stelloo et al., 2018), with the modifications described in the Supplementary Methods.

LFQ data was then further processed using *DEprot* (https://github.com/sebastian-gregoricchio/DEprot), using the built-in missForest algorithm ^51^ to impute missing data. Differential protein abundance was defined as p-value < 0.05 (paired t-test) and |LFQ difference| > 1. For RIME experiments, over-representation analysis using the WebGestalt ^52^ webtool was used against the CORUM database (v5) at default parameters for enriched proteins using the protein coding genome as a background. For whole cell proteomics the LFQ difference was used as a ranking metric before using clusterProfiler for GSEA.

### ChIP-seq experiments

For ChIP-seq experiments 42×10^6^ LNCaP cells were seeded per condition in DCC containing medium. After 72h cells were treated with 5nM R1881 or DMSO for 4h or 24h. Cells were fixed in 1% formaldehyde (Sigma-Aldrich) for 10 min and quenched with 0.125M L-glycine (Sigma-Aldrich). Cells were scraped on ice in PBS containing cOmplete EDTA-free protease inhibitor cocktail (Roche), pelleted at 2,000 rcf for 5min at 4°C and stored at -80°C until further processing. ChIP-seq and QC was performed as previously described ^53,54^ with the modifications described in the Supplementary Methods.

Differential peak analyses were performed using DiffBind ^55^ using the DESeq2 mode and applying a threshold of 1 for the |log_2_(fold-change)| and 0.05 for the false discovery rate (FDR). Signal of three biological replicates was averaged per condition using the *bigwigAverage* function from deepTools ^56^.

Annotation of peaks to genomic regions and overrepresentation analysis was done with ChIPseeker ^57^ and clusterProfiler ^58^ respectively with default parameters (TSS region = ± 3 kb). Tornado plots were generated with the *computeMatrix* and *plotHeatmap* functions from deepTools. Density plots were generated using the *plot.density.profile* function from Rseb (^59^, https://github.com/sebastian-gregoricchio/Rseb). For these, the bigwig files of QC passing replicates were merged using *bigwigAverage* from deeptools with a bin size of 50 bp. These files were also used for pyGenomeTracks from deeptools to make the tracks of a genomic region of interest. The obtained images were then adjusted in Adobe Illustrator.

GIGGLE analysis was performed using the CistromeDB toolkit (http://dbtoolkit.cistrome.org) with top 10k peaks and default parameters and the data was replotted in R.

### RNA-seq

For RNA-sequencing, cells were seeded with a density of 1×10^6^ cells (LNCaP) and 7.5×10^5^ (42D) cells in a 6cm-dish in DCC. After 72h of hormone deprivation, cells were either treated with 10nM R1881 or DMSO for 6h. RNA was isolated using the RNeasy column purification and DNAse kit (Qiagen) according to the manufacturer’s instructions. Libraries were prepared using the Illumina TruSeq polyA stranded RNA prep kit and subsequently sequenced on the NovaSeq 6000 platform. Paired-end 51 bp reads were deduplicated and quality checked before aligning to hg38 using TopHat2 ^60^. Count matrices were filtered for genes with at least 1 valid count and then used for Differential expression analysis using DEseq2 ^61^ with thresholds of adjusted p-value < 0.05 and |log_2_(fold-change)| > 1. GSEA was performed using clusterProfiler. Heatmaps were generated using the pheatmap R package ^62^ and data was previously vst normalized.

Linking of ChIP-seq and RNA-seq data was performed using Cistrome-GO ^63^ with default parameters. Regulatory potential thresholds were RP score > 0.3 and log_2_(fold-change) > 1.

### siRNA knockdown

siRNAs targeting TRIM33 (siT33-1: 5’-GUCAGUUUUCUGAUAGAC-3’ | 5’-AAUAGUGGUCUAUCAGAA-3‘, siT33-2: 5’-GGUGAAGCAUGUUAUGA-3’ | 5’-UGUGAAGUUCAUAACAUG-3’) from a previously published paper ^26^ were ordered as duplexed RNA molecules from IDT as they were too short to be ordered as DsiRNAs but the non-targeting could be ordered as a DsiRNA (siNT-I: 5’-GAACCAGCCAAGGUAGACAGUCAGA-3’ | 5’-UCUGACUGUCUACCUUGGCUGGUUCCU-3’). Additionally, a Dharmacon non-targeting (siRNA pool #5) and an ON-TARGETplus TRIM33 siRNA SMARTpool was ordered.

300,000 cells were seeded in a 6-well plate in phenol red free RPMI medium. 24h after seeding, the cells were transfected according to the RNAiMAX (Thermo Scientific) protocol and 48h later protein or RNA was harvested.

## DATA AVAILABILITY STATEMENT

Mass spectrometry data have been deposited at the ProteomeXchange Consortium through the PRIDE ^64^ partner repository with the identifier PXD058914. All sequencing data generated in this study (RNA-seq and ChIP-seq) have been deposited on GEO under accession number GSE284522.

## ACKNOWLEDGEMENTS

We thank the Zwart and Bergman group and the members of the TRIM-Net for discussion and input. The authors would like to thank the Research High Performance Computing (RHPC) facility of the NKI for the infrastructure to analyze the genomics datasets and the Genomics Core Facility (GCF) at the NKI for performing sequencing experiments.

## ADDITIONAL INFORMATION

### Author contribution

N.E. and W.Z. designed the study and planned experiments. N.E. performed the experiments together with J.J. (non LNCaP knockout data and LNCaP knockout ChIP-seq experiments), S.L. (AR RIME experiments) and N.P. (Antibody validations). Data was analyzed by N.E. with the help of S.G. and J.S.. L.H. and O.B. helped planning mass-spec experiments and ran the samples and provided initial data tables. A.B. and W.Z. supervised the experiments and analyses. N.E. and W.Z. wrote the manuscript with input from all authors.

