## Supplemental Methods and Figures for "TRIM33 loss reduces Androgen Receptor transcriptional output and H2BK120 ubiquitination"

**Supplementary Methods**

**ChIP-seq**

Cell pellets were thawed on ice, resuspended in lysis buffer 1 (LB1: 50 mM Hepes, 140 mM NaCl, 1mM EDTA, 10% glycerol, 0.5% NP40, 0.25% Triton-X100, KOH to pH 7.5; $10\times10^6$ cells/mL) and incubated on a rotator for 10 minutes at 4°C. Nuclei were pelleted at 2,000 rcf for 5 minutes at 4°C. Pellets were resuspended in lysis buffer 2 (LB2: 10 mM Tris, 200 mM NaCl, 1 mM EDTA, 0.5 mM EGTA, HCl to pH 8; $10\times10^6$ cells/mL), incubated on a rotator for 10 minutes at 4°C and centrifuged at 2,000 rcf for 5 minutes at 4°C. Washed nuclei were resuspended in lysis buffer 3 (LB3: 10 mM Tris, 100 mM NaCl, 1 mM EDTA, 0.5 mM EGTA, 0.1% Na-DOC, 0.5% lauroylsarcosine, HCl to pH 8; $30\times10^6$ cells/mL). Nuclei were sonicated for 14 cycles (30s on/off) using a PicoBioruptor (Diagenode) and chromatin was checked to be at a size around 250bp using agarose gel electrophoresis. Triton-X100 was added to the sheared chromatin at a final concentration of 1% and debris were pelleted for 12 minutes at 20,000 rcf at 4°C. The supernatant was then incubated overnight under rotation at 4°C with 50 µL (1.5 mg) of Protein A Dynabeads (Invitrogen), previously incubated with 5 µg of the respective antibody (AR, TRIM33: as above, TRIM24: NB100-2596, Novus Biologicals, H3K18ac: C15410139, Diagenode, H3K9me3: ab8898, Abcam). Beads were then washed 10 times with RIPA-ChIP buffer (50 mM HEPES, 500 mM LiCl, 1mM EDTA, 1% NP-40, 0.7% Na-DOC, pH = 7.6), washed once in TBS and reverse cross-linked at 65°C in 200 µL of Elution Buffer (EB: SDS 1%, 50mM Tris, 10mM EDTA) for 12-16 hours. Upon 30 minutes treatment with 40 µg RNase A (Life Technologies), 1 hour treatment with PK buffer (10 µL 0.5M EDTA, 20 µL 1M Tris-HCl pH 6.5, 40 µg Proteinase K (Invitrogen)), ChIP DNA was purified using phenol/chloroform/isoamylalcohol (25:24:1, pH 8.0, Thermo Scientific) and precipitated with 2 volumes of 100% ethanol. Illumina multiplex-sequencing with 51 bp paired-end setup was performed on the NovaSeq 6000 (Illumina) sequencer following manufacture instructions. All ChIP-seq samples were processed using the *SPACCa* pipeline (available at https://github.com/sebastian-gregoricchio/SPACCa) using default parameters. Briefly, FASTQ reads were mapped to the reference genome Hg38/GRCh38 using the accelerated version of the Burrows-Wheeler Aligner (BWA-MEM2 v0.5.10 [1]. Reads were filtered based on mapping quality (MAPQ ≥ 20), and duplicated reads were removed and RPGC normalized. Samples that did not pass QC were removed and all samples had at least 2 biological replicates (TRIM33 4h, TRIM33 24h).

#### Whole cell proteomics

*Mass spectrometry*

For protein digestion, frozen tissues were lysed in boiling Guanidine (GuHCl) lysisbuffer as described before [2]. Protein concentration was determined with a Pierce Coomassie (Bradford) Protein Assay Kit (Thermo Scientific), according to the manufacturer's instructions. After dilution to 2M GuHCl, aliquots corresponding to at least 1.05 mg of protein were digested twice (4h and overnight) with trypsin (Sigma-Aldrich) at 37°C, enzyme/substrate ratio 1:75. Digestion was quenched by the addition of FA (final concentration 5%), after which the peptides were desalted on a Sep-Pak C18 cartridge (Waters, Massachusetts, USA). From the eluates, aliquots were collected for proteome analysis and samples were vacuum dried and stored at -80°C until LC-MS/MS analysis.

Prior to mass spectrometry analysis, the peptides were reconstituted in 2% formic acid. Peptide mixtures were analyzed by nanoLC-MS/MS on an Orbitrap Exploris 480 Mass Spectrometer equipped with an EASY-NLC 1200 system (Thermo Scientific). Samples were directly loaded onto the analytical column (ReproSil-Pur 120 C18-AQ, 2.4 µm, 75 µm × 500 mm, packed in-house). Solvent A was 0.1% formic acid/water and solvent B was 0.1% formic acid/80% acetonitrile. Samples were eluted from the analytical column at a constant flow of 250 nl/min. For single-run proteome a 90-min gradient was employed containing a 78-min linear increase from 6 to 30% solvent B, followed by a 12-min wash.

Raw data were analyzed by DIA-NN (version 1.8) [3] without a spectral library and with "Deep learning" option enabled. The Swissprot Human database (20,395 entries, release 2022_02) was added for the library-free search. The Quantification strategy was set to Robust LC (high accuracy) and MBR option was enabled. The other settings were kept at the default values. The protein groups report from DIA-NN was used for downstream analysis in Perseus (version:

1.6.15.0) [4]. Values were Log2-transformed, after which proteins were filtered for at least 2 out of 3 valid values in at least one sample group.

#### RIME

*Mass spectrometry*

Peptide mixtures (10% of total digest) were loaded directly onto the analytical column and analyzed by nLC-MS/MS using a mass spectrometer connected to a nLC system as described in **Supplementary Table 1**. Solvent A was 0.1% formic acid/water and solvent B was 0.1% formic acid/80% acetonitrile. Peptides were eluted from the analytical column at a constant flow of 250 nL/min in a linear gradient, see **Supplementary Table 2** for gradient details. Raw data were analyzed by MaxQuant (see **Supplementary Table 2**, MaxQuant version)[5] using standard settings for label-free quantitation (LFQ). MS/MS data were searched against the Human database (see Table 2, database) complemented with a list of common contaminants and concatenated with the reversed version of all sequences. The maximum allowed mass tolerance was 4.5ppm in the main search and 0.5Da for fragment ion masses. False discovery rates for peptide and protein identification were set to 1%. Trypsin/P was chosen as cleavage specificity allowing two missed cleavages. Carbamidomethylation (C) was set as a fixed modification, while oxidation (M) and deamidation (N, Q) were used as variable modifications. LFQ intensities were Log2-transformed in Perseus (see **Supplementary Table 2**) [4], after which proteins were filtered as described in **Supplementary Table 3**.

#### Euler graphs

Euler graphs were calculated using the eulerr web tool (https://eulerr.co) and adjusted in Adobe Illustrator. Larsson J (2021). *eulerr: Area-Proportional Euler and Venn Diagrams with Ellipses*. R package version 6.1.1, https://CRAN.R-project.org/package=eulerr.

### Supplementary Figures

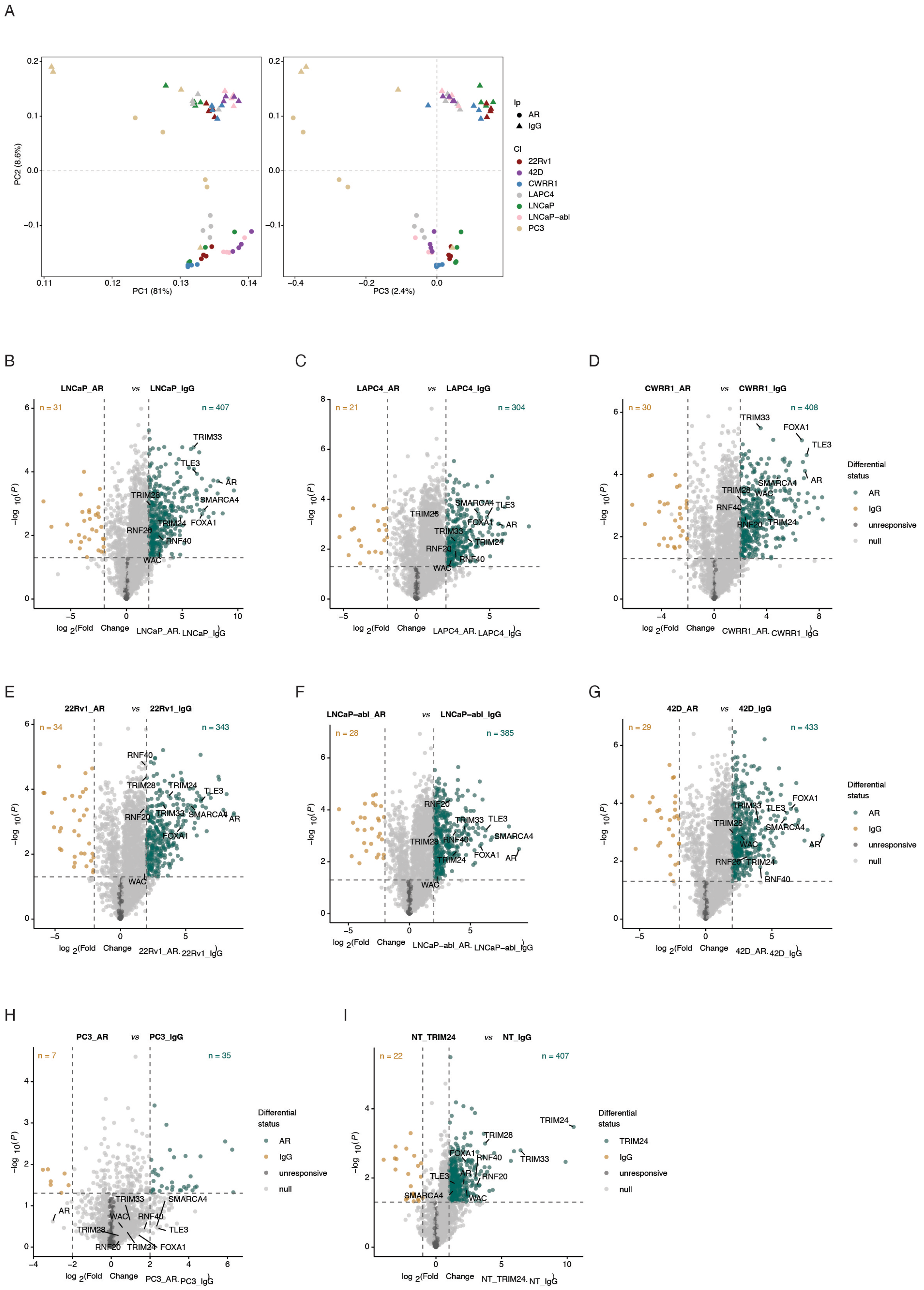


Supplementary Figure 1.

**A** PCA of all AR RIME samples including AR negative PC3 cells. **B**-**H** Volcano plots for AR across the tested cell lines. **I** Volcano plot of the TRIM24 RIME.

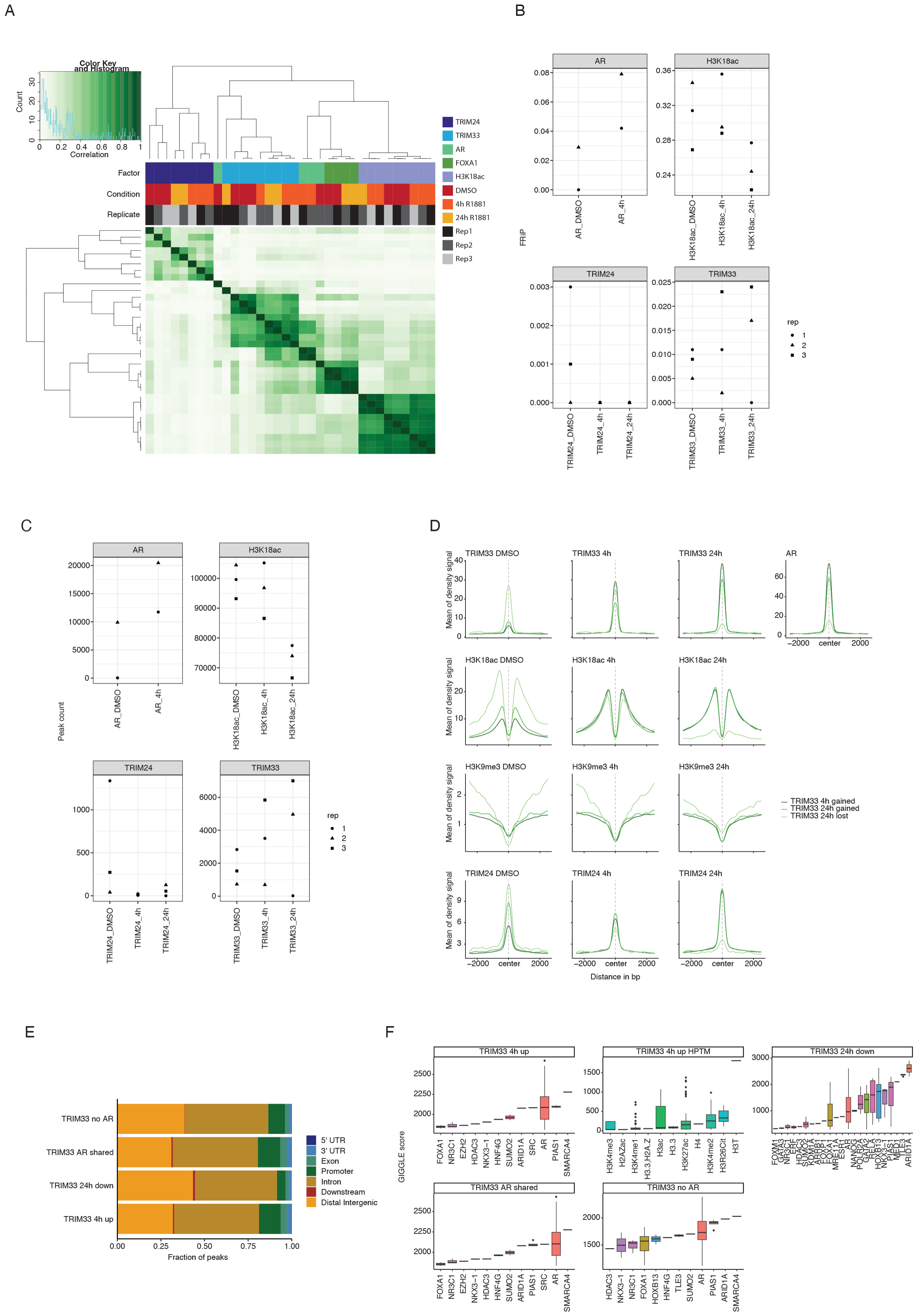

#### Supplementary Figure 2

**A** Correlation heatmap across all ChIP-seq experiments done in LNCaP wild-type cells. **B** Fraction of Reads in Peaks (FRiP) per IP and sample. AR and FOXA1 are from GSE94682. **C** Number of called peaks across samples. **D** Signal intensity plots per factor and stimulation across TRIM33 differential bound sites after stimulation (**Figure 2D**). Light green is TRIM33 24h lost, green is TRIM33 24h gained and dark green is TRIM33 4h gained. **E** Feature distribution of peak sets from **Figure 2C**. **F** GIGGLE analysis of the other subsets from **Figure 2**.

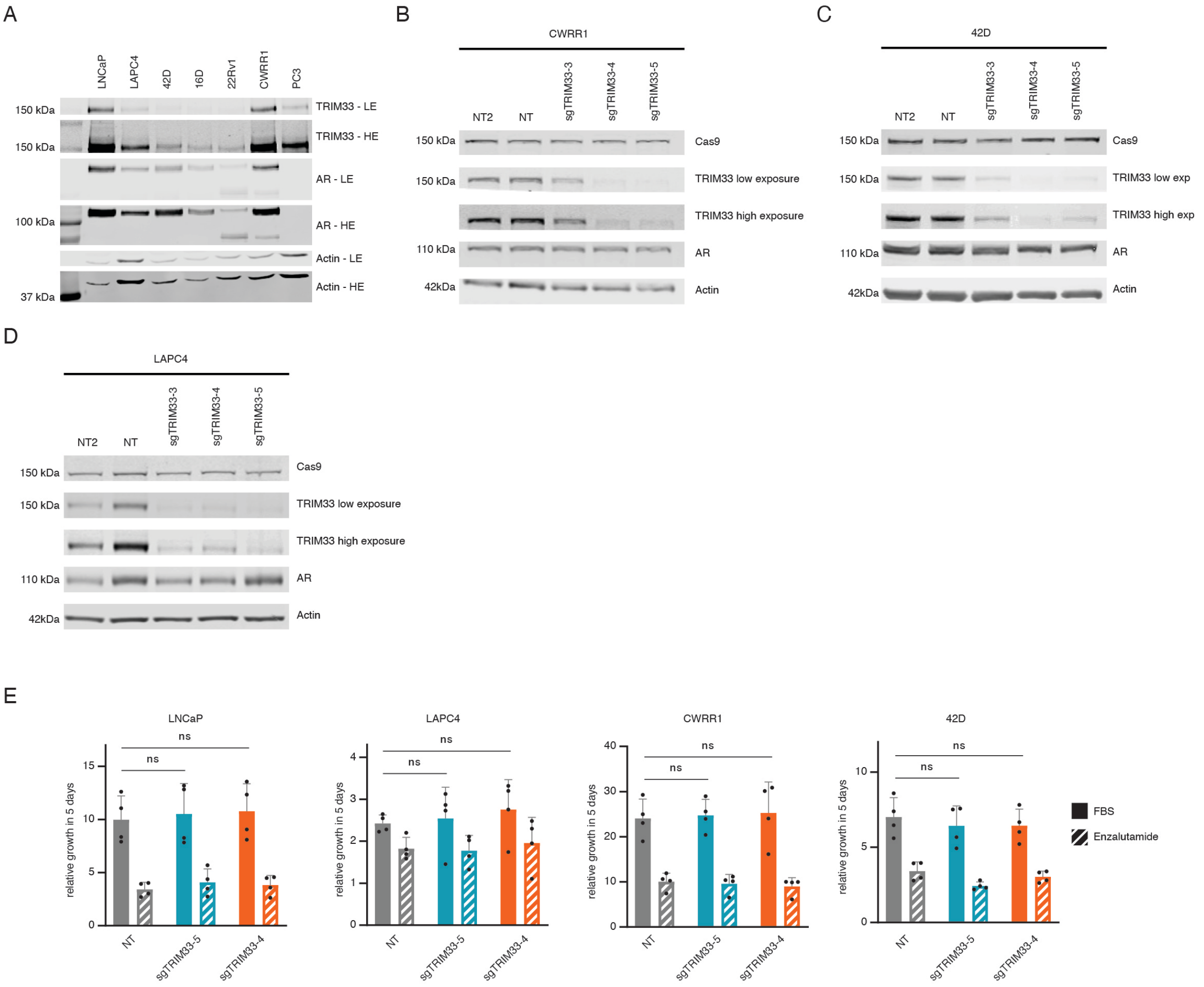


#### Supplementary Figure 3

**A** Western blot for TRIM33, AR and Actin across all used cell lines. **B-D** Western blot for TRIM33 knockout validation across several prostate cancer cell line models **E** CellTiter-Glo assay results for several polyclonal knockout cell lines with or without 10 µM Enzalutamide treatment. Error bars represent the standard deviation and statistics were calculated using a one-way ANOVA.

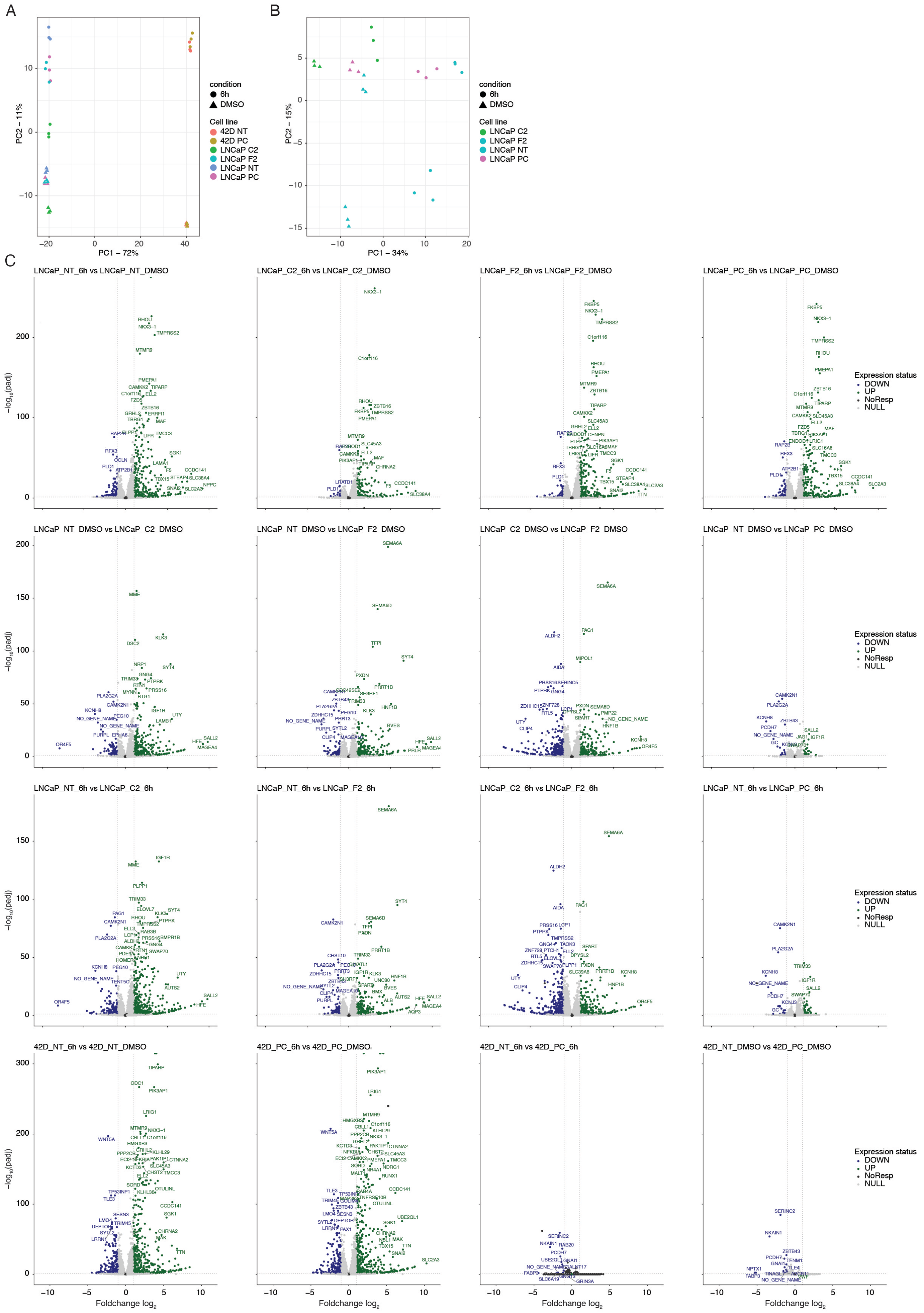


### Supplementary Figure 4

**A** PCA plot for all created RNA-seq datasets. Shapes correspond to treatment and colors to cell lines. PC = polyclonal, NT = non-targeting control. **B** PCA for only LNCaP cell lines. **C** Volcano plots for RNA-seq data of all LNCaP comparisons as well as 42D comparisons.

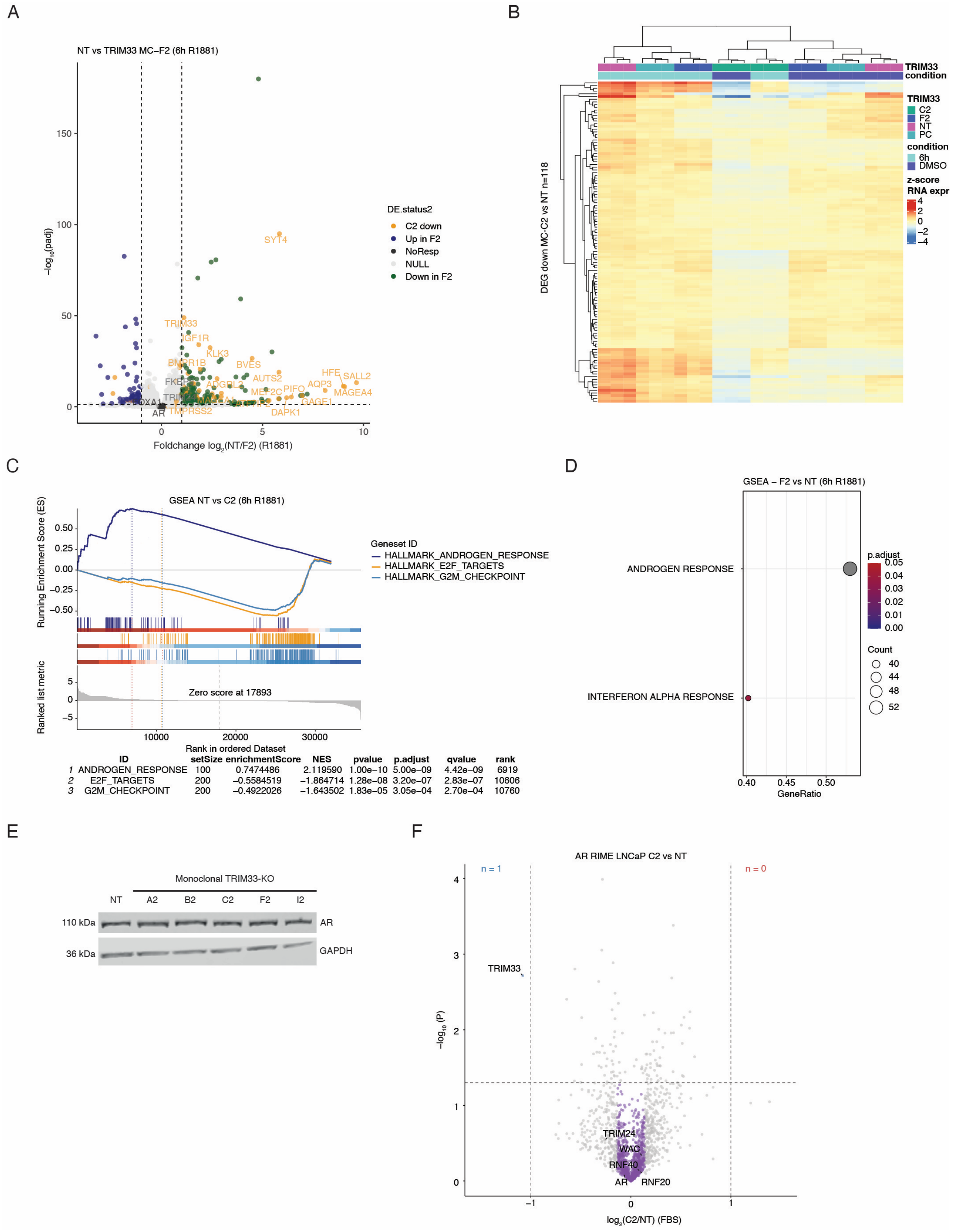


#### Supplementary Figure 5

**A** Volcano plot of TRIM33 MC-F2 versus non-targeting RNA-seq results. TRIM33 MC-C2 down genes in comparison to the non-targeting are labelled in orange. **B** Heatmap for all samples across TRIM33 MC-C2 DEG genes versus the non-targeting control. **C** Alternative representation for the GSEA results of the TRIM33 MC-C2 versus non-targeting cell line. **D**

GSEA results for TRIM33 MC-F2 versus non-targeting LNCaP cells at 6h R1881 stimulation. **E** Western blot for AR across LNCaP monoclonal knockout cell lines with GAPDH as a loading control. **F** Volcano for AR RIME in TRIM33 MC-C2 knockout cells.

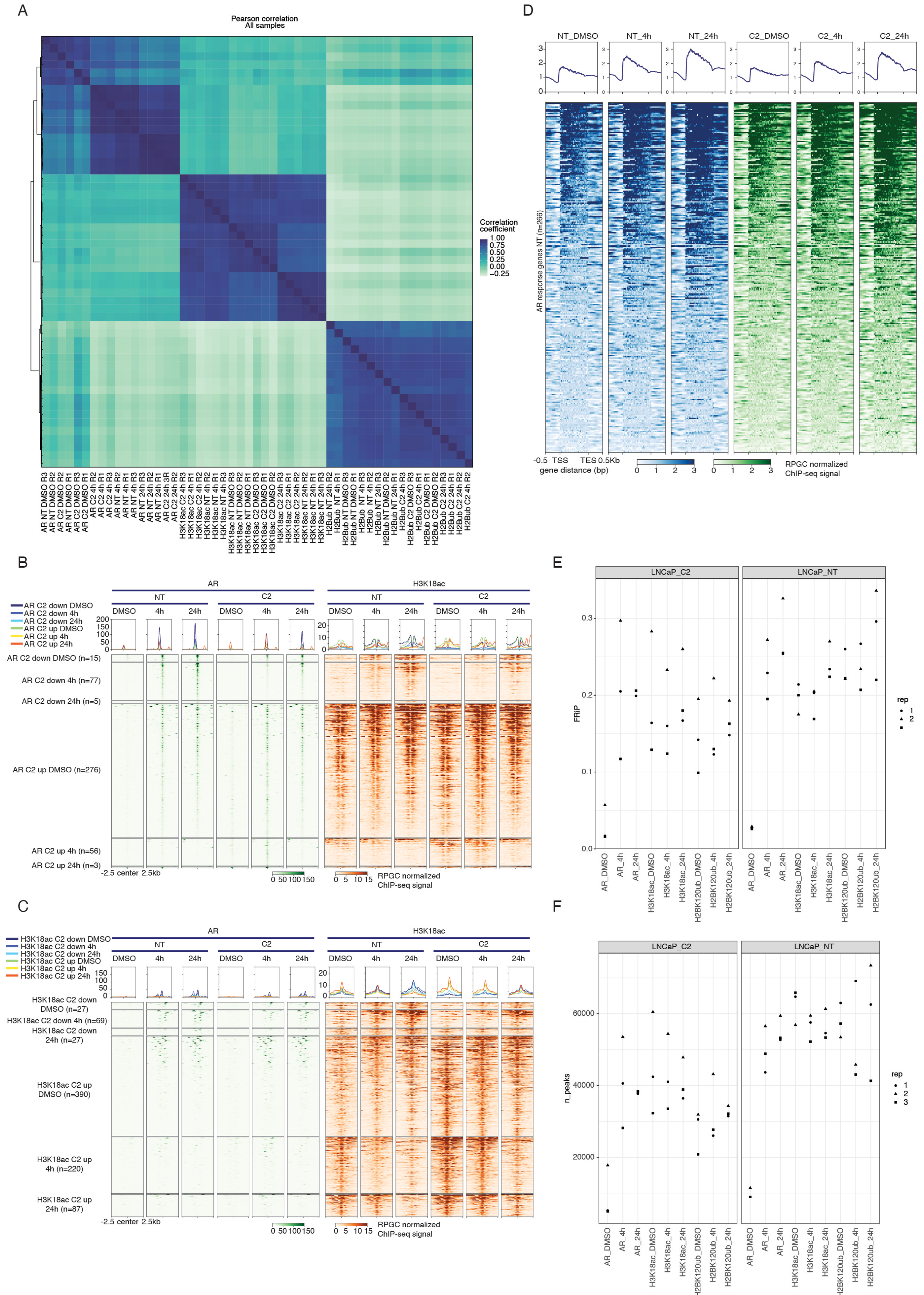

#### Supplementary Figure 6

**A** Correlation heatmap of ChIP-seq experiments in the TRIM33 MC-C2 and non-targeting cells across all IPs. **B** Heatmap for H2Bub ChIP-seq signal across samples on AR response genes. **C**

Heatmaps across differential AR peaks between the same time point (0, 4h, 24h) between C2 and NT. **D** Same as C but for H3K18ac. **E** FRiP across all samples and Ips. **F** Number of called peaks across all IPs in TRIM33 knockout and non-targeting cells.

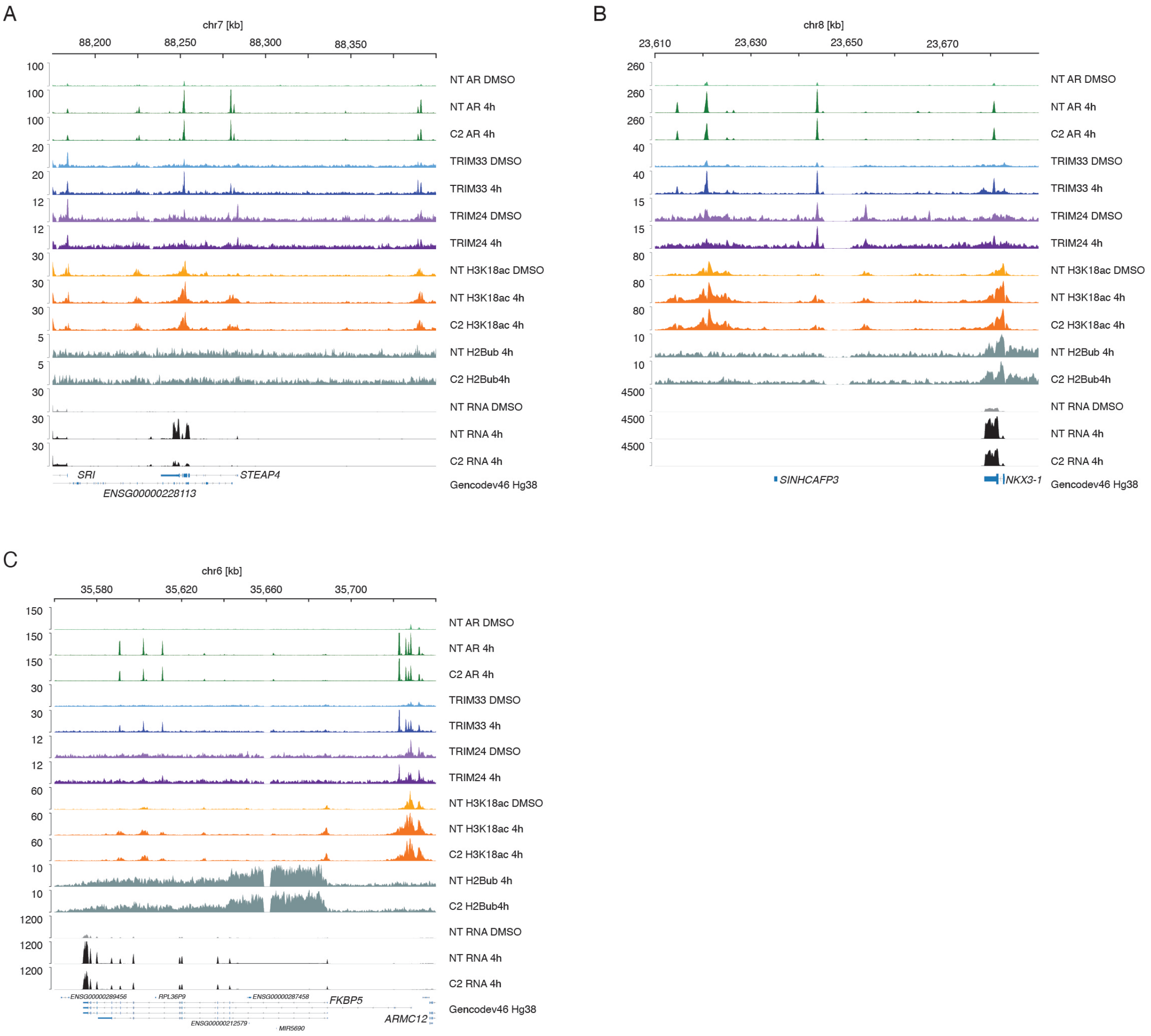


#### Supplementary Figure 7

**A** Genomic track for the *STEAP4* locus with the available ChIP-seq and RNA-seq data. Unstimulated conditions (DMSO) are displayed in lighter color than the AR activated (4h or 6h) conditions. NT represents non-targeting cells whereas C2 represents a monoclonal TRIM33 knockout. **B** Same as in A for the *NKX3-1* locus. **C** Similar to A and B for the *FKBP5* locus. The Genecode track shows all transcripts and their isoforms and is not collapsed.

A

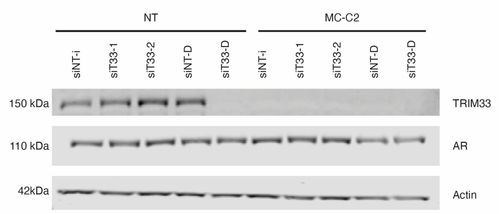


#### Supplementary Figure 8

**A** Western blot for TRIM33 and AR after siRNA transfection in LNCaP cells. siNTi = NT from Chen et al., siT33-1/2 = RNA duplex from Chen et al., siNT-D = non-targeting siRNA pool from Darmacon, siT33-D = TRIM33 targeting siRNA pool from Dharmacon. Shown is a representative of 3 replicates.
